# Establishment of Strigolactone-Producing Bacterium-Yeast Consortium

**DOI:** 10.1101/2021.06.29.450423

**Authors:** Sheng Wu, Xiaoqiang Ma, Anqi Zhou, Alex Valenzuela, Kang Zhou, Yanran Li

**Affiliations:** Department of Chemical and Environmental Engineering, University of California, Riverside, California 92521, USA; Disruptive & Sustainable Technologies for Agricultural Precision, Singapore-MIT Alliance for Research and Technology, Singapore; Department of Chemical and Biomolecular Engineering, National University of Singapore, Singapore; Department of Botany and Plant Sciences, University of California, Riverside, California 92521, USA

## Abstract

Strigolactones (SLs) are a class of phytohormones playing diverse roles in plant growth and development, yet the limited access to SLs is largely impeding SL-based foundational investigations and applications. Here, we developed *Escherichia coli*-*Saccharomyces cerevisiae* consortia to establish a microbial biosynthetic platform for the synthesis of various SLs, including carlactone, carlactonic acid, 5-deoxystrigol (5DS, 6.65±1.71 µg/L), 4-deoxyorobanchol (4DO, 3.46±0.28 µg/L), and orobanchol (OB, 19.36±5.20 µg/L). The SL-producing platform enabled us to conduct functional identification of CYP722Cs from various plants as either OB or 5DS synthase. It also allowed us to quantitatively compare known variants of plant SL biosynthetic enzymes in the microbial system. The titer of 5DS was further enhanced through pathway engineering to 47.3 µg/L. This work provides a unique platform for investigating SL biosynthesis and evolution and lays the foundation for developing SL microbial production process.

## Introduction

Strigolactones (SLs) were initially characterized as signaling molecules, which are released from plant roots, induce germination of root parasitic weed, regulate the hyphae branching of arbuscular mycorrhiza fungi (AMF), and promote the symbiotic relationship between plants and fungi (*1, 2*). Later, they were also identified as a novel class of plant hormones that control shoot branching, leaf growth and senescence, and promote the formation of lateral root and growth of primary root (*3*). SLs thus have been considered as promising agrochemicals, such as bio-stimulants that enhance the nutrient uptake efficiency through modulating plant-AMF symbiotic association (*4-6*). To date, more than 30 natural SLs have been isolated (fig. S1) (*7, 8*). SLs generally consist of a conserved butenolide ring (D ring) connected to a less conserved tricyclic lactone ring via an enol-ether bond (Fig. 1) (*9*). They can be classified into canonical and non-canonical SLs: the canonical SLs contained the tricyclic lactone-ring (ABC ring), while the non-canonical SLs lack of the tricyclic ring scaffold with one (C ring) or two rings (B ring and C ring) missing (*10*). The canonical SLs can be further subdivided into orobanchol (*O*)- and strigol (*S*)-type SLs according to the stereochemistry in the C ring, which are represented by 4-deoxyorobanchol (4DO) and 5-deoxystrigol (5DS), respectively (*9*). Some of the better-known non-canonical SLs include methyl carlactonoate (MeCLA) (*11*), heliolactone (*12*), avenaol (*13*), zealactone (*14*), and lotuslactone (*15*).

**Fig. 1.**
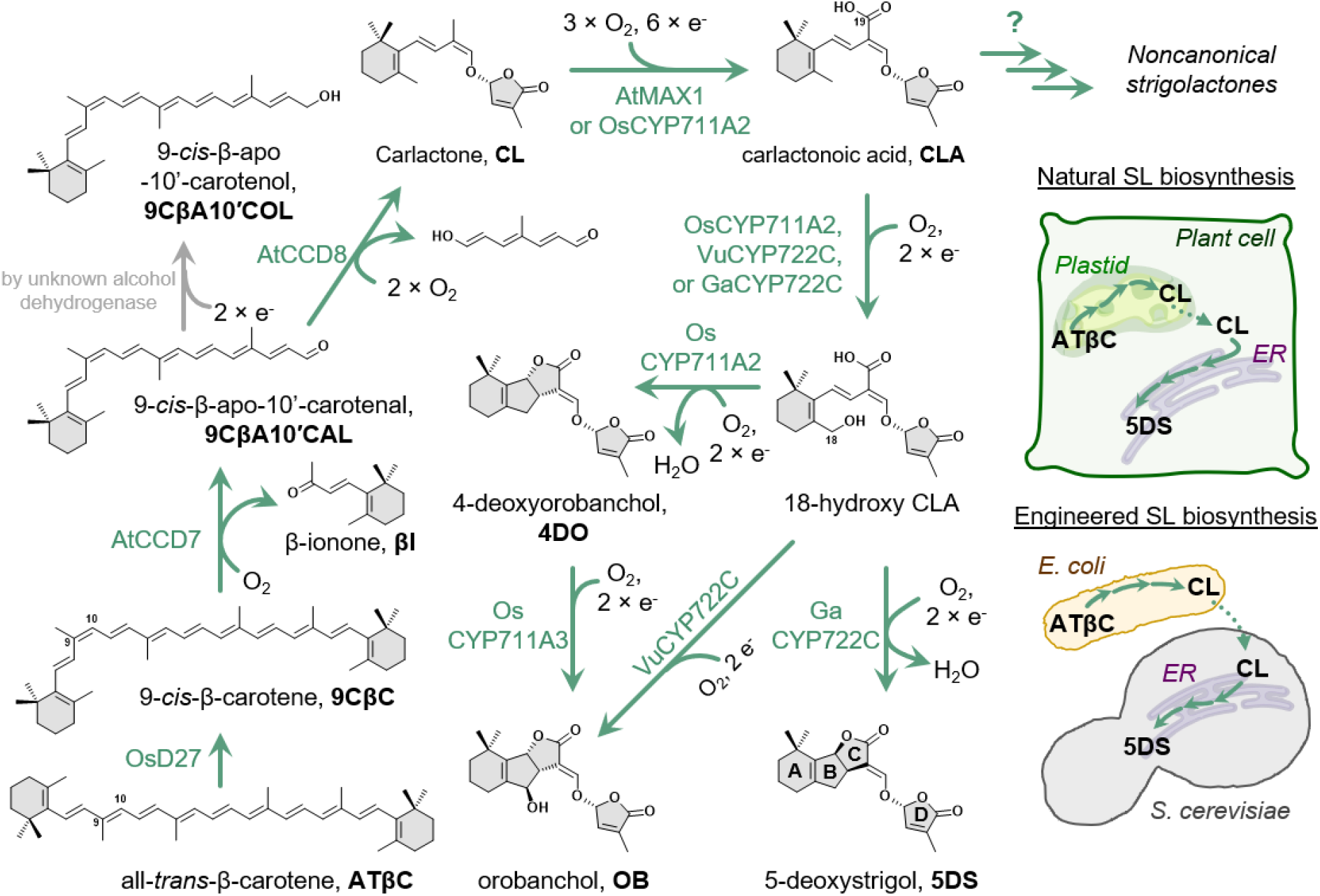
Mimicking plant strigolactone pathway distribution in the engineered *E. coli*-*S. cerevisiae* co-culture. OsD27, DWARF27 from *Oryza sativa*; AtCCD7, carotenoid cleavage dioxygenase 7 from *Arabidopsis thaliana*; AtCCD8, carotenoid cleavage dioxygenase 8 from *Arabidopsis thaliana*; AtMAX1, MORE AXILLARY GROWTH 1 from *Arabidopsis thaliana*; OsCYP711A2, cytochrome P450 CYP711A2 from *Oryza sativa*; OsCYP711A3, cytochrome P450 CYP711A3 from *Oryza sativa*; VuCYP722C, cytochrome P450 CYP722C from *Vigna unguiculata*; GaCYP722C, cytochrome P450 CYP722C from *Gossypium arboreum*; Carlactone is an important branch point. NADPH-cytochrome P450 reductase 1 from *Arabidopsis thaliana* (ATR1) is expressed in yeast for the functional reconstitution of the plant cytochrome P450s. ER: endoplasmic reticulum.

SLs are derived from all-*trans*-β-carotene (ATβC), which is converted to carlactone (CL), the key branching point in SL biosynthesis (*16*), by the functions of three plastid enzymes: the isomerase DWARF27 (D27), and carotenoid cleavage dioxygenases 7 and 8 (CCD7 and CCD8) (Fig. 1) (*16*). D27, a [2Fe-2S]-containing polypeptide, catalyzes the isomerization of ATβC to 9-*cis*-β-carotene (9CβC) (*16, 17*), followed by CCD7, a non-heme iron-dependent enzyme that catalyzes the C9′-C10′ double bond cleavage of 9CβC to yield 9-*cis*-β-apo-10′-carotenal (9CβA10′CAL) and β-ionone (βI) (*16, 18, 19*). Subsequently, CCD8 further catalyzes the oxidative cleavage of 9CβA10′CAL to synthesize CL, with the reaction mechanism remaining elusive (*16, 20*). CL is then exported into cytoplasm, and further oxidized by cytochrome P450s (CYPs) and other oxidases to afford various SL structures (*21*). The first oxidation step has been characterized to be the C19-oxidation of CL to synthesize carlactonic acid (CLA), which can be catalyzed by the MORE AXILLARY GROWTH 1 (MAX1), a member of the CYP711A subfamily (*11*).

MAX1 is conserved in a number of plant species (*22*) and has many homologs. Rice, known to produce *O*-type SL such as 4DO and orobanchol (OB) (*23*), encodes five MAX1 homologs. One MAX1 homolog, OsCYP711A2 encoded by *Os900*, was identified to catalyze the conversion of CL to 4DO likely *via* CLA (*22, 24*). The B ring of 4DO can be further oxidized by MAX1 analogs (OsCYP711A3 encoded by *Os1400* from rice, ZmMAX1b from maize) to afford OB (*22*). However, 4DO is not produced in many OB-producing plants (e.g., cowpea), which hints another synthetic route of OB without involving 4DO (*25, 26*). This direct synthesis of OB from CLA was later identified to be catalyzed by CYP722C in cowpea (*27*). Recently, a CYP722C from cotton was characterized to catalyze the synthesis of 5DS from CLA (*28*).

Biosynthesis of most other SLs have not been fully elucidated. Studying these pathways could be sped up if total biosynthesis of key SLs and pathway intermediates (e.g., CL, CLA, OB, 4DO and 5DS) from sugar can be established in fast-growing microbial hosts. *In situ* produced SLs and pathway intermediates in the microbial system can serve as substrates to characterize recombinant SL biosynthetic enzyme candidates, eliminating the need of acquiring these molecules from plant materials or chemical synthesis. On the other hand, much remains to be investigated to understand the evolutionary history of SL biosynthesis and signaling in plants, especially non-vascular plants. The SL biosynthetic genes are either missing from the non-vascular plants or quite distinct from the corresponding variants encoded by the vascular plants (*29*). For example, although Charales (green algae) have been reported to produce SLs (e.g., sorgolactone) (*30*), no close CCD7 and CCD8 analogs have been found from either expression profiles or genomes, whether Charales can synthesize SL or not remains in debate (*29*). An efficient functional identification strategy aided by a microbial SL-producing platform can also advance understanding of SL origin and evolution.

In this study, we first attempted to establish the SL biosynthetic pathway in *Saccharomyces cerevisiae* but failed, possibly due to the unsuccessful functional reconstitution of D27 and CCD7. We then tested the use of *Escherichia coli* as the host, since the active form of D27, CCD7, and CCD8 have been heterologous expressed and isolated from *E. coli* before (*16, 18-20, 31*). An engineered *E. coli* strain successfully produced CL but failed to further functionalize it. The challenge was later overcome by using mixed cultures of *E. coli* and *S. cerevisiae*. Total biosynthesis of OB, 4DO, and 5DS was achieved from xylose. We utilized this microbial biosynthetic platform to identify eight CYP722Cs which produced either OB or 5DS from CLA and established a sequence-function correlation that could be used to predict the function of unknown CYP722Cs. Next, we chose 5DS as a model SL and improved the titer in the microbial consortium by 220% to 47.3 µg/L (shake flask culture) through metabolic engineering. The improved titer could be useful to isolate adequate pathway intermediates in future studies. Finally, we quantitatively compared variants of D27, CCD7 and CCD8 from different plant species in the optimized system.

## Results

### It is challenging to establish CL production in *S. cerevisiae*

To synthesize CL, we first attempted to functionally reconstitute D27 and CCD7 in a ATβC-producing *S. cerevisiae* strain. We reconstructed the ATβC-producing yeast strain (**YYL23**, table S2) as previously described (*32*). Trace amount of 9CβC naturally existed in the established ATβC-producing yeast strain with a ratio to the ATβC approximately 1:25 (fig. S2A). Expressing D27 from *A. thaliana* (AtD27) or rice (OsD27) did not increase the ratio (fig. S2A). We extensively explored commonly used strategies for functionally reconstituting D27 (protein N-terminal engineering, protein localization to mitochondria, improving [2Fe-2S] cluster biogenesis, and reducing oxidative stress; fig. S2, A to C), but without success. We also tried to reconstitute the function of CCD7 from *A. thaliana*, due to the presence of the substrate of CCD7, 9CβC, though at a low titer in **YYL23**. Although CCD1 has been functionally expressed in yeast to cleave β-carotene to afford the synthesis of βI (*33*), we did not detect any activity of CCD7 possibly due to the fact that the CCD1 is natively localized in cytosol while the other CCDs are in plastid (fig. S2D) (*34*). Truncating N-terminus of CCD7 and/or targeting it to yeast mitochondria, which resemble plastids to certain extent, did not lead to detection of CCD7 activity in the yeast (fig. S2D).

### Establishment of CL production in *E. coli*

Previous investigations indicate that D27, CCD7, and CCD8 can be expressed and isolated in soluble form from *E. coli* for the *in vitro* biochemical investigations (*16, 18-20, 31*). Thus, we shifted the CL production from yeast to *E. coli*. First, OsD27 was expressed from a medium-copy number plasmid *pCDFDuet-1* (table S1) in *E. coli*, under the control of *T7* promoter, in the presence of the well-documented ATβC-producing plasmid *pAC-BETAipi* (*35*) (resulting strain **ECL-2**, *E. coli* harboring *pAC-BETAipi* to produce ATβC is designated as **ECL-1**, table S2). Upon the introduction of OsD27, the ratio between 9CβC to ATβC was increased from 1:4.1 to 1:1.3, which indicates the functional reconstitution of OsD27 in *E. coli* (Fig. 2A and fig. S3, A and B). In the presence of OsD27, the titer of 9CβC was 1.41 mg/L. The activity of D27 from *A. thaliana* (AtD27) was also examined in **ECL-1** (**ECL-*2″***, table S2), which exhibited a lower activity than OsD27 (fig. S4). We truncated the putative plastid transit peptide (first 40 amino acids) from OsD27 but did not detect obvious enhancement in the conversion towards 9CβC (fig. S4). Previous investigation on D27 indicated that this β-carotene isomerase catalyzes reversible conversion between ATβC and 9CβC (*31*), and thus the ratio between 9CβC to ATβC in **ECL-2** might have reached the equilibrium.

**Fig. 2.**
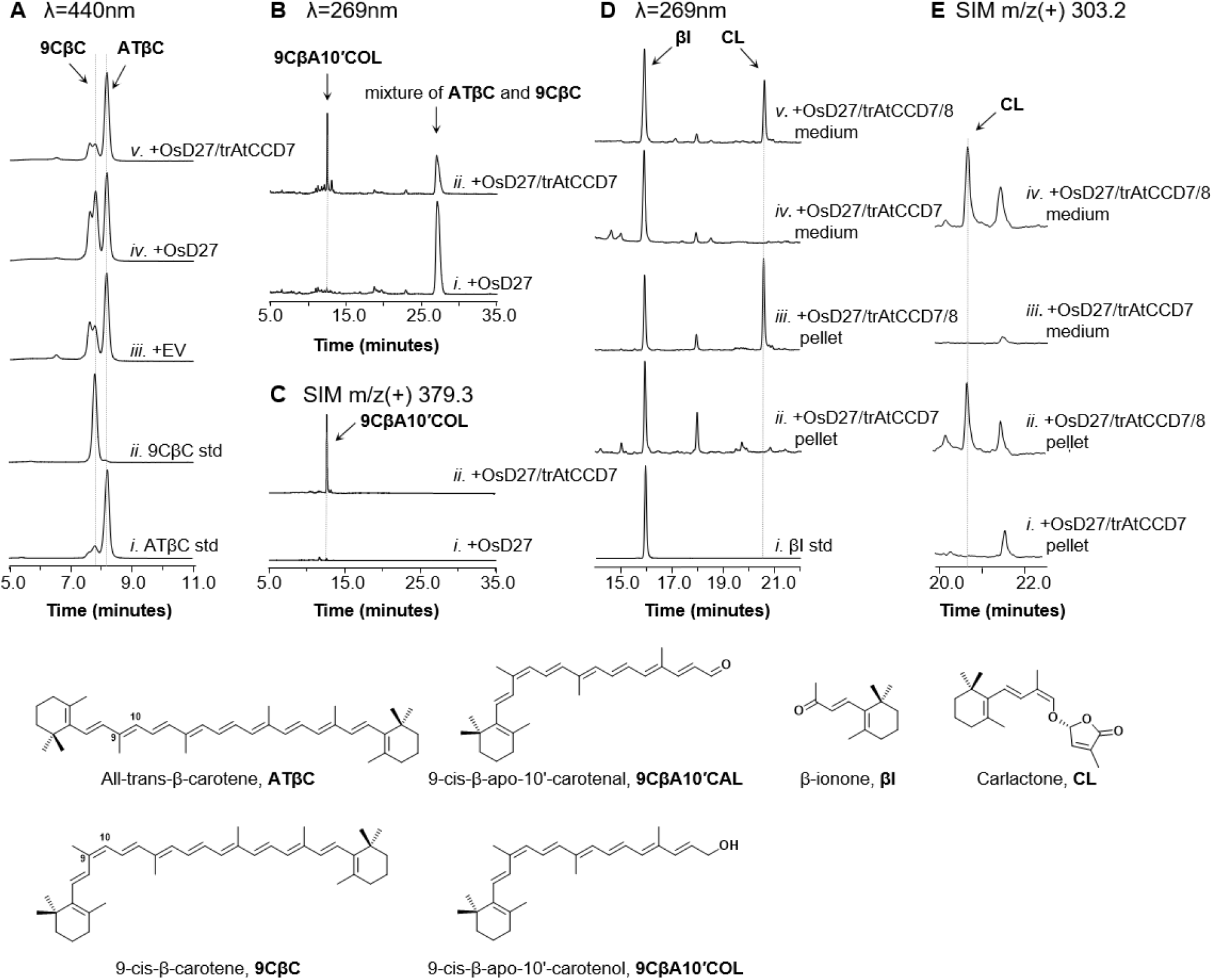
Production of CL in *E. coli*. Information of all the microbial strains mentioned in the caption (strain names are in bold font) can be found in table S2. (**A**) HPLC analysis (λ=440 nm) of i) ATβC standard, ii) 9CβC standard, cell extracts of *E. coli* harboring iii) ATβC-producing plasmid (*pAC-BETAipi*, table S1) and empty plasmid (EV) (**ECL-1**), iv) *pAC-BETAipi* and OsD27-expressing plasmid (**ECL-2**), v) *pAC-BETAipi* and OsD27/trAtCCD7 co-expressing plasmid (**ECL-3**). The samples were analyzed using *Separation Method I* (see Materials and Methods). (**B**) HPLC analysis (λ=390 nm) of cell extracts of i) **ECL-2**, ii) **ECL-3** by using *Separation Method II* (see Materials and Methods). (**C**) Selected ion monitoring SIM (SIM) extracted ion chromatogram (EIC) at 9CβA10′COL’s characteristic m/z^+^=379.3 of cell extracts of i) **ECL-2**, ii) **ECL-3**. *Separation Method II* was used. (**D**) HPLC analysis (λ=269 nm) of i) βI standard, 9CβA10′COL-producing *E. coli* harboring ii) EV (cell pellet extracts, **ECL-4N**), iii) trAtCCD8-expressing plasmid (cell pellet extracts, **ECL-4**), iv) **ECL-4N** (medium extracts), v) **ECL-4** (medium extracts) using *Separation Method III* (see Materials and Methods). (**E**) SIM EIC at CL’s characteristic m/z^+^ signal (MW=302.19Da, [C_19_H_26_O_3_+H]^+^=[C_19_H_27_O_3_]^+^=303.2) of i) **ECL-4N** (cell pellet extracts), ii) **ECL-4** (cell pellet extracts), iii) **ECL-4N** (medium extracts), iv) **ECL-4** (medium extracts). *Separation Method III* was used. All traces are representative of at least three biological replicates for each engineered *E. coli* strain.

Subsequently, the N-terminus of CCD7 from *A. thaliana* (AtCCD7) was truncated (first 31 residues (*31*), trAtCCD7) and introduced to 9CβC-producing **ECL-2** from the same plasmid *pCDFDuet-1* expressing OsD27, under the control of *T7* promoter (**ECL-3**, table S2). Liquid chromatography-mass spectrometry (LC-MS) analysis indicated the synthesis of a new compound with m/z^+^=379.3 that agrees with 9-*cis*-β-apo-10′-carotenol (9CβA10′COL) (Fig. 2, B and C, and fig. S3C) (*16, 18, 19*). Although the product of CCD7 *in vitro* is 9CβA10′CAL (*16*), the aldehyde can be reduced to the corresponding alcohol, 9CβA10′COL, in *E. coli* (*18, 19*). The synthesis of βI, the other product of CCD7, was confirmed through comparison with the authentic standard (figs. S3D and S5). No 9CβA10′COL (Fig. 2C) but a trace amount of βI was detected in **ECL-2** (fig. S5), and βI is believed to be produced by the autoxidation of ATβC (*36*).

The N-terminal truncated (first 56 residues) AtCCD8 (trAtCCD8) was then expressed in the 9CβA10′COL-producing **ECL-3** from a medium-copy number plasmid *pET21a* under the control of *T7* promoter (**ECL-4**, table S2). Although a drastic decrease in the synthesis of 9CβA10′COL was detected upon the introduction of trAtCCD8, we were unable to detect the synthesis of CL (fig. S6, A and B). Previous reports imply that CL was unstable (*31, 37*), and that pH, solvent composition, and temperature can affect its chemical stability (*37, 38*). We added 200 mM HEPES buffer (pH 7.0) to the growth medium upon IPTG induction (buffer volume / medium volume = 0.5), which led to the detection of a tiny new peak in chromatograms of both the cell pellets and the culture medium at RT=20.58 min with m/z^+^=303.2, which agrees with that of CL (figs. S3E and S6, C and D). However, the yield of CL was low (fig. S6, C and D), and was not proportional to the decrease in the synthesis of 9CβA10′COL. Thus, we tested different medium conditions for the detection of CL. Under the optimal fermentation conditions (XY medium; fig. S6, E to G), a distinguished peak with the maximum absorption at 269 nm and m/z^+^ consistent with those of CL at the same retention time as the putative CL was observed (Fig. 2, D and E, and fig. S3E) (*16, 31*).

### Synthesis of CLA in *E. coli*-*S. cerevisiae* coculture

According to the pioneering *in planta* study, most of the canonical SLs are branched from CLA, which is synthesized from CL with the function of MAX1 (*11*). Although *E. coli* does not contain membrane-bound organelles, many plant cytochrome P450s have been functionally reconstitute in *E. coli (39)*. However, further introduction of truncated MAX1 and cytochrome P450 reductase from *A. thaliana* (*40*) (AtMAX1 and ATR1, respectively) in the CL-producing **ECL-4** did not convert CL towards CLA (**ECL-5**, table S2, fig. S8). Considering AtMAX1 is an endoplasmic reticulum (ER)-localized enzyme, the eukaryotic model species *S. cerevisiae* maybe a more suitable organism to reconstitute the activity of the ER-localized CYP than *E. coli*(*39*). Since CL can be detected in both cell pellets and culture medium, it is possible to establish the synthesis of downstream SLs using an *E. coli*-*S. cerevisiae* coculture, with which CL is expected to be translocated from *E. coli* to yeast for further functionalization.

AtMAX1 and ATR1 were then introduced to *S. cerevisiae* on two low-copy number plasmids and expressed downstream of *PGK1* and *TEF1* promoter, respectively (new strain: **YSL-1**, table S2). When the CL-producing *E. coli* strain (**ECL-4**) was co-cultured with **YSL-1**, the peak of CL substantially decreased, and a new compound was detected in the organic extract of both cell pellets and medium under UV detection (Fig. 3A). The UV-VIS spectrum of the new peak had the maximum absorption at 271 nm, with [M-H]^-^ = 331.1 matching those of CLA (Fig. 3B and fig. S3F). HRMS analysis confirmed [M-H]^-^ = 331.1627 consistent with that of CLA (C_19_H_24_O_5_; fig. S7A). We then tried to improve the CLA titer by adjusting the inoculum ratio between **ECL-4** and **YSL-1**. We found that increasing the amount of yeast led to increased conversion of CL towards CLA (fig. S9). When the *S. cerevisiae*-*E. coli* inoculum ratio increased from 0.2 to 2, the ratio between CLA to CL rose from 0.3 to 17.3. When the *E. coli*-yeast inoculum ratio was higher than 1, CLA production was not further improved, so we used the ratio in the subsequent experiments.

**Fig. 3.**
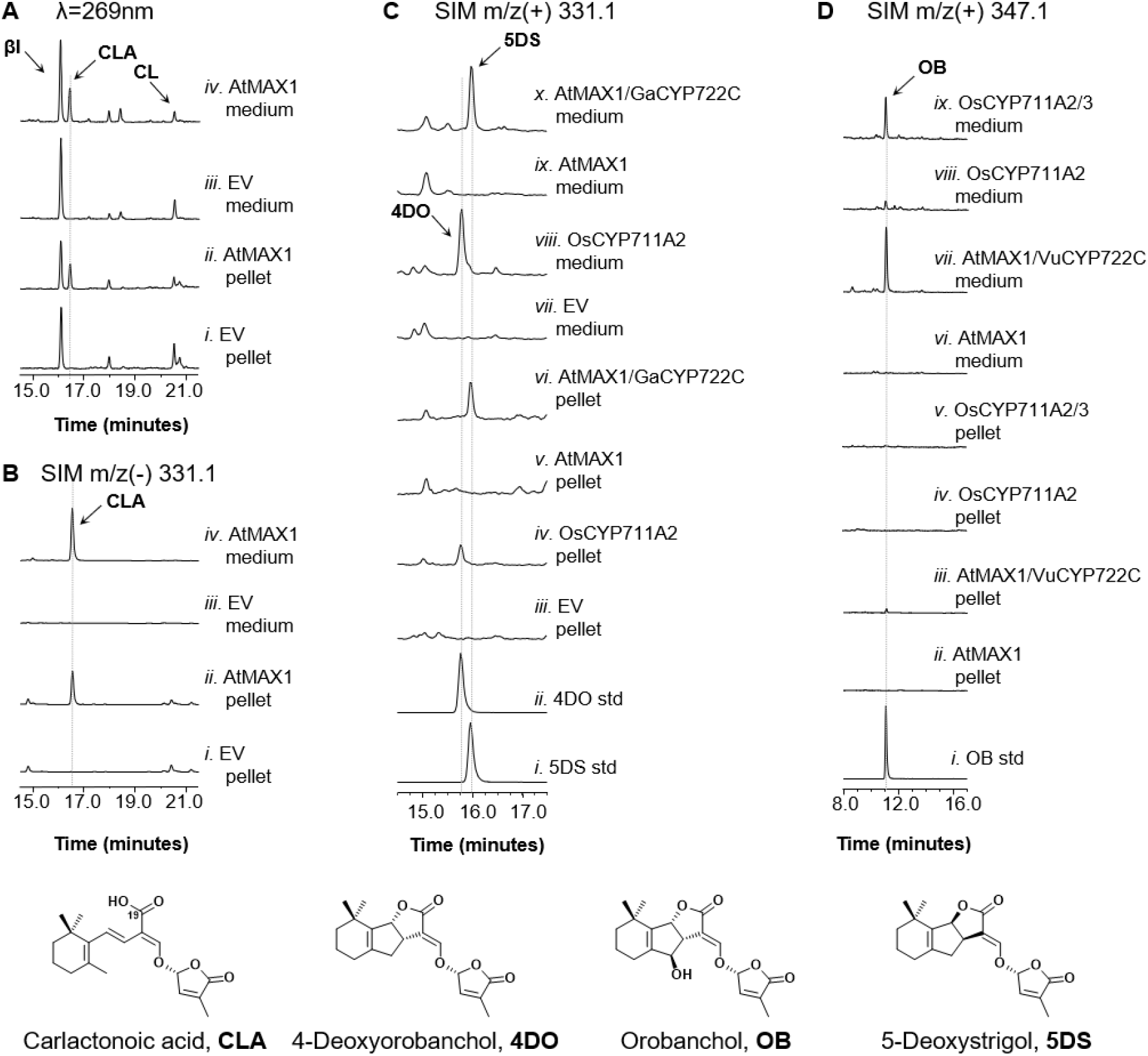
Production of CLA and three canonical strigolactones using *E. coli*-*S. cerevisiae* co-cultures. *Separation Method III* was used for all the chromatograms in this figure. Information of all the microbial strains mentioned in the caption (strain names are in bold font) can be found in table S2. (**A**) HPLC analysis (λ=269 nm) of CLA in CL-accumulating *E. coli* (**ECL-4**) cocultured with ATR1-expressing *S. cerevisiae* harboring i) empty vector (EV, cell pellet extracts, **YSL-1N**), ii) AtMAX1-expressing plasmid (cell pellet extracts, **YSL-1**), iii) **YSL-1N** (medium extracts), iv) **YSL-1** (medium extracts). (**B**) SIM EIC using CLA’s characteristic m/z^-^ signal (MW=332.16Da, [C_19_H_24_O_5_-H]^-^=[C_19_H_23_O_4_]^-^=331.1) of CL-producing **ECL-4** cocultured with i) **YSL-1N** (cell pellet extracts), ii) **YSL-1** (cell pellet extracts), iii) **YSL-1N** (medium extracts), iv) **YSL-1** (medium extracts). (**C**) SIM EIC using 4DO or 5DS’s characteristic m/z^+^ signal (MW=330.15Da, [C_19_H_22_O_5_+H]^+^=[C_19_H_23_O_5_]^+^=331.1) of i) 5DS standard; ii) 4DO standard; cell pellet extracts of CL-accumulating **ECL-4** cocultured with ATR1-expressing *S. cerevisiae* harboring iii) empty vector (EV, **YSL-2N**), iv) OsCYP711A2-expressing plasmid (**YSL-2**), v) AtMAX1-expressing plasmid (**YSL-4/5N**), vi) AtMAX1/GaCYP722C-expressing plasmids (**YSL-5**); and medium extracts of **ECL-4** co-cultured with vii) **YSL-2N**, viii) **YSL-2**, ix) **YSL-4/5N**, x) **YSL-5**. (**D**) SIM EIC using OB’s characteristic m/z^+^ signal (MW=346.14Da, [C_19_H_22_O_6_+H]^+^=[C_19_H_23_O_6_]^+^=347.1) of i) OB standard; cell pellet extracts of CL-accumulating **ECL-4** cocultured with ATR1-expressing *S. cerevisiae* harboring ii) AtMAX1-expressing plasmid (**YSL-4/5N**), iii) AtMAX1/VuCYP722C-expressing plasmids (**YSL-4**), iv) OsCYP711A2-expressing plasmid (**YSL-3N**), v) OsCYP711A2/OsCYP711A3-expressing plasmids (**YSL-3**); medium extract of **ECL-4** cocultured with vi) **YSL-4/5N**, vii) **YSL-4**, viii) **YSL-3N**, ix) **YSL-3**. All traces are representative of at least three biological replicates for each engineered *E. coli*-*S. cerevisiae* consortium.

### Synthesis of OB through 4DO using CYP711As in *E. coli*-*S. cerevisiae* coculture

In rice, the two MAX1 homologs, OsCYP711A2 and OsCYP711A3, were identified to catalyze the conversion of CL to 4DO and 4DO to OB, respectively (*22, 24*). To synthesize 4DO in the microbial system, we expressed *Os900* (OsCYP711A2 gene) using a low-copy number plasmid downstream of *PGK1* promoter in a yeast strain harboring ATR1 (**YSL-2**, table S2). When **ECL-4** was co-cultured with **YSL-2**, a new peak with m/z^+^ at 331.1 (consistent with either 5DS or 4DO) was detected (Fig. 3C), which was further confirmed to be 4DO through comparison with the authentic 4DO and 5DS standards (Fig. 3C and figs. S3G, and S10, A to C). The titer of 4DO in the consortium was 3.46±0.28 µg/L. In addition to 4DO, we were also able to detect the synthesis of another new compound in comparison to the negative control (without OsCYP711A2) in the organic extract of the medium. The putative new compound showed m/z^-^ at 347.1 in the negative-ion mode (figs. S3H and S10B), and is putatively 18-hydroxy-CLA (*27*). HRMS analysis confirmed that [M-H]^-^ = 347.1521 and the fragmentation pattern are both consistent with those of 18-hydroxy-CLA reported previously (C_19_H_24_O_6_; fig. S7B) (*27*).

Furthermore, we introduced *Os1400* (OsCYP711A3 gene) to **YSL-2** on a low-copy plasmid driven by *GPD* promoter (**YSL-3**, table S2). As expected, compared to the **ECL-4**/**YSL-3N** (**YSL-3N** is equivalent to **YSL2** but harboring an additional empty vector; table S2) coculture, the **ECL-4**/**YSL-3** coculture synthesized substantially less 4DO and 0.75±0.01 µg/L OB (identified and quantified by using the authentic OB standard; Fig. 3D and figs. S3I and S11). Different from 4DO detected in both pellets and medium, OB mostly in the medium indicates that most OB synthesized was exported into the medium. The successful reconstitution of OB synthesis from CL in the microbial consortium using OsCYP711A2 and OsCYP711A3 confirmed the previously proposed synthetic pathway of OB in rice (*24*).

### Synthesis of OB using CYP722C in *E. coli*-*S. cerevisiae* coculture

Previous investigations confirmed that VuCYP722C from cowpea can directly convert CLA into OB and its diastereomer ent-2′-epi-orobanchol by *in vitro* experiments (*27*). To establish this OB biosynthetic route in the microbial system, we introduced VuCYP722C to **YSL-1** on a low-copy number plasmid downstream of the *GPD* promoter (**YSL-4**, table S2), then co-cultured **YSL-4** with the CL-producing **ECL-4**. The **ECL-4**/**YSL-4** co-culture produced much less CLA than the **ECL-4**/**YSL-4/5N** co-culture (**YSL-4/5N** is equivalent to **YSL1** but harboring an additional empty vector; table S2, fig. S12A), and synthesized 19.36±5.20 µg/L OB (Fig. 3D and fig. S12C). Interestingly, we were also able to detect the putative 18-hydroxy-CLA in the medium of the **ECL-4**/**YSL-4** co-culture (fig. S12B). Our results confirmed that OB can be generated through two different routes using different set of CYPs.

### Synthesis of 5DS using CYP722C in *E. coli*-*S. cerevisiae* coculture

GaCYP722C from cotton was reported to be responsible for the conversion of CLA into 5DS through an *in vitro* investigation (*28*). To synthesize 5DS in microbial consortium, we introduced the *GaCYP722C* gene into the CLA-producing yeast on a low-copy number plasmid downstream of *GPD* promoter (**YSL-5**, table S2). When the new yeast strain was co-cultured with the CL-producing **ECL-4**, we detected 6.65±1.71 µg/L of 5DS (5DS was identified and quantified by using the authentic standard; Fig. 3C and figs. S3J and S13C). Different from OB, 5DS was detected in the extract of both medium and cell pellet, which algins with the lower hydrophilicity of 5DS than OB (fig. S13). Only a small quantity of CLA was detected (fig. S13A), suggesting the high efficiency of GaCYP722C. The putative 18-hydroxy-CLA was also detected in the medium of the 5DS-producing consortium (fig. S13B). This peak exists in the medium of all the canonical SL-producing consortiums we constructed, which supports the hypothesis that 18-hydroxy-CLA is a common intermediate in canonical SLs biosynthesis.

### Functional Mapping of Various CYP722Cs

CYP722C genes are widely distributed in flowering plants. GaCYP722C share 65% amino acid identity with VuCYP722C, yet they catalyze different reactions. It is intriguing to investigate whether the enzymatic function of homologous proteins is conserved across different plant species. The successful functional reconstitutions of GaCYP722C and VuCYP722C in the microbial consortium hints the potential of using this biosynthetic platform to establish a sequence-function correlation of CYP722Cs, which will enable predicting the function of unknown CYP722Cs. We used GaCYP722C protein sequence as a query, performed BLASTp search, and selected a total of 28 CYP722C sequences from different plant species including dicotyledons and monocotyledons (table S4). SL profile of some of these plant species have been examined. For example, birdsfoot trefoil (*Lotus japonicus*) (*2*) and woodland strawberry (*Fragaria vesca*) were reported to produce 5DS (*37*), while cowpea (*Vigna unguiculata*) (*26, 41*), red bell pepper (*Capsicum annuum*) (*25*), and red clover (*Trifolium pratense*) were reported to produce OB without involving 4DO (*25, 41*). Tomato (*Solanum lycopersicum*) produces OB (*42*). We also included two CYP722Cs with the functions to be confirmed (SlCYP722C (*27*) and LjCYP722C (*43*)), and CYP722A and CYP722B sequences as the outgroup (Fig. 4).

**Fig. 4.**
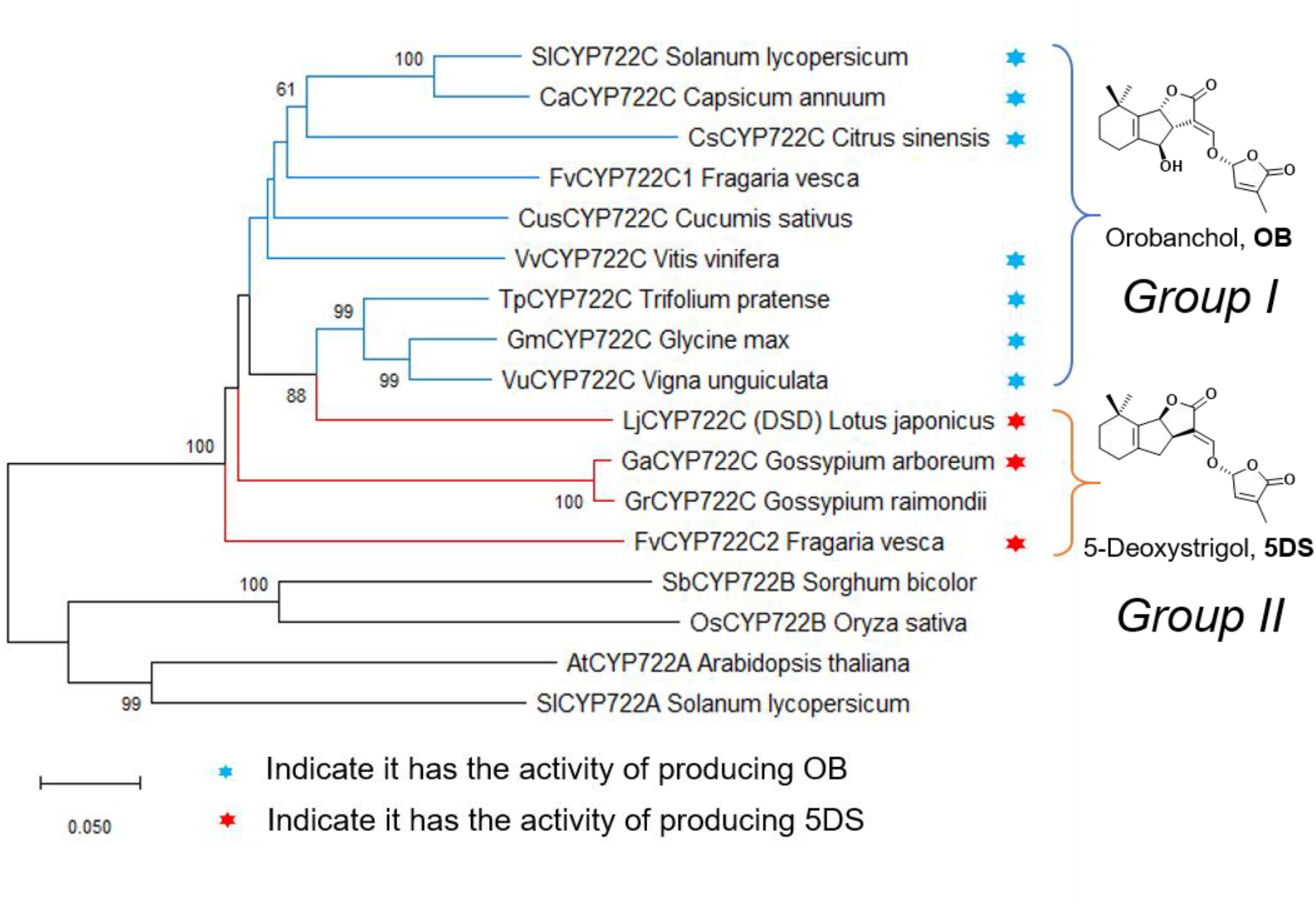
Phylogenetic analysis of CYP722C and the functional mapping. Phylogenetic analyses were conducted in MEGA X by using ClustalW for multiple sequence alignment and the neighbor-joining method. The parameters are set as follows, bootstrap test (1000 replicates), p-distance mode, complete deletion. The numbers next to branches indicate bootstrap values (>60% are shown). This analysis involved 17 amino acid sequences, including CYP722A and CYP722B sequences to root the tree. The accession numbers of proteins are listed in table S4. The asterisk means that these enzymes have been tested in this study, blue asterisk indicates OB-producing activity and red asterisk indicates 5DS producing activity.

Phylogenetic analysis indicated that CYP722C subfamily can be divided into two groups (Fig. 4 and fig. S15): Group I and Group II. The characterized OB-producing VuCYP722C and SlCYP722C are members of Group I. The speculative 5DS synthase LjCYP722C and characterized 5DS-producing GaCYP722C are members of Group II. To examine if we can simply predict the function based on the phylogenetic analysis, we synthesized eight CYP722C genes (table S3) from different branches and screened their functions by introducing each gene to the CLA-producing microbial consortium on a low-copy number plasmid regulated by *GPD* promoter (table S2). Among the eight CYP722C genes we examined, all the CYP722Cs from Group I (*S. lycopersicum, C. annuum, T. pratense, Glycine max* [Soybean], *Citrus sinensis* [Sweet orange], and *Vitis vinifera* [Grape]) indeed converted CLA to OB (Fig. 4 and fig. S16A), while the two CYP722Cs from Group II (*F. vesca* and *L. japonicus*) synthesized 5DS from CLA (Fig. 4 and fig. S16B). We also examined the function of OsCYP722B (*O. sativa*) and SbCYP722B (*Sorghum bicolor*) using the CLA-producing consortium and did not detect any conversion of CLA or synthesis of 5DS or OB (table S2 and fig. S14). The CYP722C characterization results are consistent with the previously reported SL profiles from the corresponding plants (*2, 25, 26, 37, 41, 42*), and suggest that sweet orange and grape are capable of synthesizing OB though their SL profiles have not been reported.

The SL-profile of most plants is not reported. The phylogenic analyses and functional characterization of CYP722Cs from various plant species in the microbial system imply a sequence-function correlation, which can be used to predict the SL synthetic capacity of the corresponding plants. If a plant encodes a group I CYP722C, likely it is able to produce *O*-type SLs; while if a group II CYP722C is present, this plant should own the capability to synthesize *S*-type SLs.

### Metabolic engineering to improve 5DS production

After demonstrating microbial biosynthesis of various SLs, we used metabolic engineering to improve product titer, which should facilitate future gene characterization efforts. We chose 5DS as the model SL. We first improved the CL-producing *E. coli* (**ECL-4**). In this strain, the building blocks of CL (isopentenyl diphosphate [IPP] and dimethylallyl diphosphate [DMAPP]) were supplied by an endogenous, non-engineered methylerythritol phosphate (MEP) pathway. Biosynthesis of many isoprenoids was found in prior studies to be limited by this pathway (*44-47*), possibly due to that *E. coli* evolved to regulate this pathway to have low flux due to weak demand of isoprenoids by the cell. To upregulate the MEP pathway, we overexpressed its first enzyme, 1-deoxy-D-xylulose-5-phosphate synthase (EcDXS, coded by *dxs*). Since *dxs* is in the same operon in the *E. coli* genome with *ispA* (a gene also involved in the CL biosynthetic pathway), the dxs-ispA operon including its native promoter and terminator was inserted into *pAC-BETAipi*. IspA encodes farnesyl diphosphate synthase (EcEPPS) and would also be upregulated in the new strain (**ECL-6**, table S2). When **ECL-4** was replaced by **ECL-6** in the 5DS-producing co-culture (**ECL4**/**YSL-5**), the pool size of CL was increased by 150% (Fig. 5A), but the 5DS titer was not increased (Fig. 5B). Since the increase in the CL pool size could also be due to reduced consumption of CL, we analyzed a by-product (βI) of the CL biosynthetic pathway. Because there was no reaction known to consume βI in this culture, its titer can be used as an indicator of the CL production rate. The βI titer of the **ECL-6**/**YSL-5** co-culture was 330% (Fig. 5C) higher than that of the **ECL-4**/**YSL-5** co-culture, suggesting that the overexpression of *dxs* and *ispA* indeed improved the CL production in the co-culture, and that it is needed to improve the conversion of CL into 5DS in **YSL-5**.

**Fig. 5.**
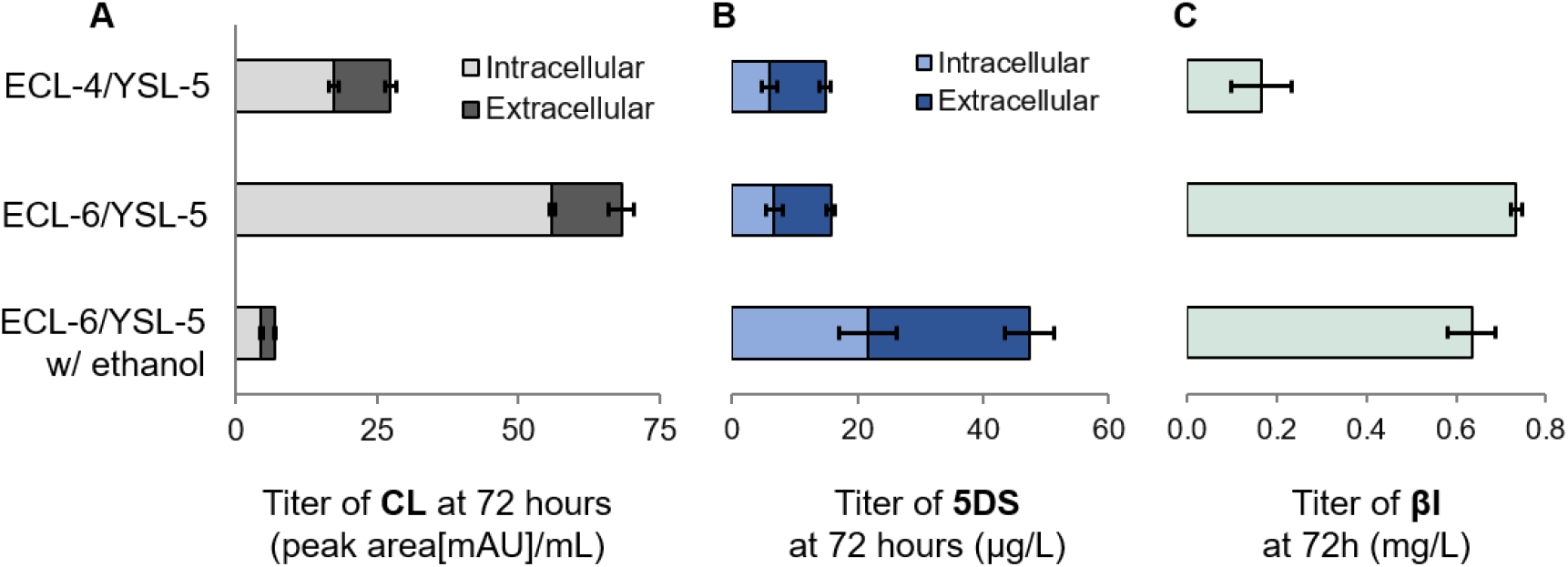
Pathway engineering and process optimization improved 5-deoxystrigol (5DS) titer. Information of all the microbial strains mentioned in the caption (strain names are in bold font) can be found in table S2. The native *ispA-dxs* operon was introduced to **ECL-4** (resulting strain: **ECL-6**) to overexpress *E. coli* 1-deoxy-D-xylulose-5-phosphate synthase (EcDXS) and farnesyl diphosphate synthase (EcFPPS). At 72 h after starting the co-culture, we measured 5DS titer of (**A**) CL, (**B**) 5DS, and (**C**) βI of **ECL-4/YSL-5** coculture, **ECL-6/YSL-5** coculture with and without ethanol feeding (ethanol was fed at 48 and 60 h to increase ethanol concentration in the medium by 1 g/L). All the cocultures were performed at 22°C. 5DS and βI were extracted and quantified by using authentic standards as described in Materials and Methods. Due to the lack of the authentic standard, the quantities of CL were estimated using raw data (the peak area from chromatogram of absorbance at 269 nm, RT: 31.72 min, and the unit of peak area is mAU*min and was defined as arbitral unit [a.u.]). The error bars represent the standard error of three biological replicates.

Since the SL biosynthesis is less understood, it was more difficult to pinpoint the rate-limiting step. We then decided to increase the concentration of all the enzymes involved in functionalizing CL by enlarging the population of **YSL-5**. The yeast growth was limited by acetate supply (acetate concentration constantly below detection limit, 0.1 g/L, during the co-culture; acetate was the main carbon source of the yeast and was produced by *E. coli*), so ethanol was added as a supplement carbon source for *S. cerevisiae*. Ethanol can be converted into acetate in the yeast and cannot be metabolized by *E. coli (48)*. The yeast growth was indeed improved by supplementing 4 g/L ethanol (fig. S17), and almost no CL accumulated under such condition (Fig. 5A). More importantly, 5DS titer was increased by 220% (to 47.3 μg/L; Fig. 5B) compared with the co-culture without the ethanol addition.

In addition, OsD27 in **ESL-6** was replaced with six D27 variants from different plant species individually (*Oryza nivara, Oryza rufipogon, Oryza meridionalis, Triticum aestivum, Zea mays, Populus trichocarpa*). Although none of these replacements further increased the 5DS production (fig. S18), these exercises quantitatively compare the activities of variants of SL biosynthetic enzymes from different plant species, which may shed light on evolution of SL biosynthesis (elaborated below).

## Discussion

The biosynthetic pathway from ATβC to CL is conserved in most land plants (*49, 50*), however, the routes for the conversion of CL into canonical SLs can be distinct among different plant species, even when they produce the same SLs (*50*). A microbial platform of producing CL, CLA, and the downstream pathway intermediates may make the characterization of the unknown pathways much easier. Our study indicates that it is highly challenging to establish the biosynthesis of CL in yeast, but possible in *E. coli*. Through using the *E. coli*-yeast coculture strategy, we established the biosynthesis of CL in *E. coli* and then various SLs in the *E. coli*-yeast consortium. The SL-producing microbial consortium, although currently with low SL titer (at the level of tens of µg/L), provides a handy platform for the functional characterization of SL biosynthetic enzymes, as in the proof-of-concept experiment on CYP722Cs (fig. S16). Through simply introducing the CYPs of interest to the yeast strain of the corresponding SL-producing microbial consortium (for CYP722Cs, CLA-producing consortium), we were able to identify the function of CYP722Cs within two weeks upon arrival of the synthetic genes. This approach does not need to supply synthetic CLA, which is chemically unstable and commercially unavailable. The functions of the CYP722Cs identified by the microbial system were consistent with the reported SL profiles of the corresponding plants.

SLs exhibit extremely low abundance in nature (up to 70 pg of OB/plant detected in the roots of red clover seedlings) (*51, 52*), making isolation of SLs from plant materials to be laborious and costly. Due to the structural complexity and instability of SLs, the chemical synthesis of SLs is laborious and expensive to be an economic SL supply strategy for the SL-based agricultural applications (*4, 9, 53*). Chemical synthesis has been useful in SL-related research through providing some synthetic SL analogues (*4*), such as GR24. However, synthetic analogs are generally less active than natural SLs (*54*) and generally in racemic mixtures, while different isomers exhibit different activities (*55*). For example, the commonly used synthetic SL analog GR24 is 100-fold less potent at stimulation of *Orobanche minor* germination than natural SLs (*56*), and (+)-GR24 is 100 times more active than (-)-GR24 as a branching factor (*57*). Unfortunately, many conclusions on the biological effects of SLs are obtained using racemic GR24 (*6, 8, 58*). In addition, SLs of subtle structural differences may exhibit different bioactivities (*49, 59*). For example, different SLs have been found to exhibit distinct efficiencies towards shoot branching inhibition in plants and hyphal branching activity in arbuscular mycorrhizal (AM) fungi (*49*). The limited access to natural SLs is hindering comprehensive investigations on the structure-activity correlation of this group of phytohormones in plant. The development of efficient microbial bioproduction of various natural SLs will advance such investigations and agricultural applications.

Much remains to be investigated into the biochemistry and evolution of CYP722Cs. Consistent with the pioneering *in planta* and biochemical characterizations of CYP722Cs (*27, 28*), our study also implies that CYP722Cs are divided into two major groups, one group converts CLA to OB (Group I) and the other one synthesizes 5DS (Group II). Both types of CYP722Cs are believed to synthesize OB or 5DS through first catalyzing 18-hydroxylation of CLA (*27, 28*). Group I CYP722C further catalyzes the oxidation of the alcohol at C18 to aldehyde that triggers nucleophilic attack to afford B and C ring closure for the synthesis of OB (the oxygen atom is retained at C-18). This process would not produce 4DO (fig. S19) (*27*). On the other hand, Group II CYP722C does not catalyze further oxidation of 18-hydroxyl; it may protonate the 18-hydroxyl to serve as a leaving group to afford the synthesis of 5DS (fig. S19) (*28*). Interestingly, in the functional identification of various CYP722Cs, we did not detect the synthesis of 4DO in the CYP722C-expressing SL-producing microbial consortium. Further investigations into CYP722Cs from other plants may help answer whether there is a 4DO-synthesizing CYP722C. It is also interesting to understand how CYP722Cs evolve to control the C18-oxidation and stereospecificity of ring-closing.

A compound detected in the 4DO-producing microbial consortium (**ECL-4**/**YSL-2**) may putatively be 18-hydroxy-CLA (fig. S10B). The synthesis of 4DO by OsCYP711A2 in **ECL-4**/**YSL-2** is likely through a similar mechanism as GaCYP722C except that OsCYP711A2 catalyzes one more oxidation at C19 (to carboxylic acid) in addition to the C18-oxidation (to alcohol; fig. S19) (*24*). We did not detect any 5DS or OB in the 4DO-producing microbial consortium using OsCYP711A2 (**ECL-4**/**YSL-2**). Further biochemical investigations are needed to understand how CYP711As that convert CL to CLA were evolved to gain C18-hydroxylation ability and to catalyze the B- and C-ring closure with strict stereospecificity. The microbial platform we have built should be able to expediate enzyme characterizations to answer these questions.

In addition, the establishment of deoxy-SL (4DO and 5DS) producing microbial consortium may provide a handy platform for the discovery of the downstream tailoring enzymes in SL biosynthesis. Hydroxylation on SLs can alter the biological activities. For example, *Striga*-susceptible maize cultivar only produces 5DS, while mainly sorgomol is detected in the resistant cultivar (*60*); Sorghum produces both 5DS and sorgomol, and can convert 5DS to sorgomol (*26*). Recently, CYP728B35 encoded by Sorghum has been suggested to be the sorgomol synthase catalyzing the hydroxylation of 5DS to afford the synthesis of sorgomol (*61*). In addition to Sorghum, strigol-producing cotton converts 5DS to strigol and strigyl acetate (*26*). The enzymes involved in these oxidations of SLs remain to be characterized.

Moreover, the biosynthesis of non-canonical SLs remains largely unknown. In *Arabidopsis*, CLA can be methylated to form methylcarlactonoate (MeCLA) by an unknown methyltransferase (*11*). Recently, lateral branching oxidoreductase (LBO), a 2-oxoglutarate and Fe^II^-dependent dioxygenase, was suggested to catalyze the hydroxylation of MeCLA to hydroxymethyl carlactonoate (1’-HO-MeCLA) (*62, 63*), which may be the precursors of many non-canonical SLs such as avenaol (*37*). In addition to MeCLA and 1’-HO-MeCLA, diverse non-canonical SLs, including hydroxyl CL (3-HO-CL, 4-HO-CL, 16-HO-CL) and the corresponding hydroxyl CLA derivatives, were also detected in *Arabidopsis (63)*, with little known about the enzymes involved in the formation of these structures. *L. japonicus* can produce non-canonical SL lotuslactone (*15*), and a 2-oxoglutarate-dependent dioxygenase (2-OGD) (named *LLD*) gene was proposed to be involved in the biosynthesis of lotuslactone (*43*), yet the catalytic function of LLD was not characterized. The CL/CLA-producing microbial consortium may also provide a convenient way to functionally identify and discover the enzymatic mechanism in the biosynthesis of various SL structures, both canonical and non-canonical.

Another intriguing yet mysterious question to be answered is the origin and evolutionary history of SL biosynthesis and perception. SLs have been reported to be present in several bryophytes and green algae (*29, 30, 64*), which do not always encode the full set of SL biosynthetic enzymes as described above (*29, 30*). These facts spur debate about the origin and evolution of SL biosynthesis and perception (*29, 30*). The moss *Physcomitrella patens* encodes one set of D27, CCD7, and CCD8, but no MAX1 analog, which is consistent with previous investigation that *P. patens* only synthesizes CL (*29, 37*). The activities of *P. patens* CCD7 (PpCCD7) and CCD8 (PpCCD8) towards the synthesis of CL have been validated through *in vitro* reconstitution (*65*). On the other hand, CCD8 and MAX1 analogs are absent from the genome of liverworts *Marchantia polymorpha* (*29, 66*). In this study, the activity of *P. patens* D27 (PpD27), PpCCD8, and *M. polymorpha* CCD7 (MpCCD7) were examined in the CL-producing **ECL-6** through replacing the corresponding ortholog (e.g., PpD27 replacing OsD27, fig. S20). Replacing OsD27 with PpD27 retained the synthesis of CL, while replacing AtCCD7 or AtCCD8 with MpCCD7 or PpCCD8 completely abolished CL production (fig. S20). The inconsistency in the activity of PpCCD8 between our study and previous investigations might be due to different N-terminus truncation or codon optimization method (we truncated four more amino acids to be consistent to trAtCCD8, table S6). MpCCD7 does not encode a chloroplast transfer peptide and is more than 10% longer in length than regular CCD7s, and the activity of MpCCD7 might be different from regular CCD7s and requires further investigations (e.g., assay with more substrates than 9CβC). Furthermore, since MAX1 and CYP722 analogs are universally present in flowering plants yet generally absent from primitive plants (*29, 67*), we also examined the activity of some MAX1 and CYP722 analogs from *Sphagnum fallax* (moss, SfMAX1, 46.1 % identity to AtMAX1; SfCYP722C, 36.2 % identity to GaCYP722C) and *Klebsormidium nitens* (green algae, four MAX1 orthologues, KnMAX1a-1d, 27.9-31.2% identity to AtMAX1) in the CL-producing consortium (*66*). Unfortunately, none of these CYPs examined converted CL to CLA or CLA to 5DS or OB in the microbial consortia (fig. S21), suggesting different functions from AtMAX1 or Group I or II CYP722Cs. These species may produce SL-like compounds that has not been discovered yet, with distinct biosynthetic pathway from seed plants, which need to be characterized. Comprehensive investigations into the putative SL biosynthetic enzymes from primitive plants are necessary to fully elucidate the SL biosynthesis from these plants as well as origin and evolution of SL biosynthesis.

The current titer of 5DS (∼50 µg/L) needs to be further improved in order to isolate pathway intermediates of SL biosynthesis for structural verification (e.g., 18-hydroxy-CLA) and mechanistic investigation of SL biosynthetic enzymes (e.g., CYP722Cs). We engineered an *E. coli* strain that could produce ∼200 mg/L ATβC by using the recently developed isopentenol utilization pathway (IUP) (*68*). But when we introduced the CL-synthesizing pathway into this strain, much less CL was produced compared with using the engineered MEP pathway. A future study should investigate how to efficiently convert ATβC into CL in high ATβC-producing strains. A special attention should be paid to D27, which is the first enzyme in this segment and may control its flux. D27s from different species could be screened and the best natural D27 could be further improved by directed evolution.

Furthermore, although our study demonstrates that microbial consortium is a promising mechanism for the supply of various SLs, single-strain-based SL-producing microorganism is still desired due to the easier manipulation during fermentation. To reconstruct SL-producing *E. coli* strains, further engineering on the functional reconstitution of multiple CYPs (i.e., CYP722Cs or CYP711As) in *E. coli* is critical. On the other hand, mechanistic investigation into the failed reconstitution of D27 and CCDs in yeast may enable the rational design of microenvironment to functionally reconstitute these plastid-localized enzymes in yeast.

As a conclusion, in this study, we have successfully reconstituted the biosynthesis of CL, CLA, and three canonical SLs (4DO, 5DS, OB) in *E. coli*-yeast microbial consortia. Our study highlights the usefulness of the microbial platform as a convenient and rapid way to characterize the function of SL biosynthetic enzymes.

## Materials and Methods

### Chemicals and general culture conditions

(±)5-deoxy-strigol (purity >98%) and (±)-orobanchol were purchased from Strigolab (Italy). (±)4-deoxyorobanchol (also named as (±)-2’-epi-5-deoxystrigol) were acquired from Chempep Inc. (USA). βI is purchased from Fisher Scientific (USA). ATβC and 9CβC were purchased from Sigma-Aldrich Co. (USA). The chemically competent *E. coli* strain TOP10 (Life Technologies) was used for DNA manipulation and amplification, and was grown at 37 °C in lysogeny broth (LB) medium (Fisher Scientific) supplemented with appropriate amount of antibiotics (100 μg/mL ampicillin [Fisher Scientific], 50 μg/mL kanamycin [Fisher Scientific], 25 μg/mL chloramphenicol [Fisher Scientific], and/or 50 μg/mL spectinomycin [Sigma-Aldrich]) for plasmid maintenance. For protein expression and CL-production, we used chemically competent *E. coli* strain BL21(DE3) (Novagen), and LB (or XY medium). XY medium contains 13.3 g/L KH_2_PO_4_, 4 g/L (NH_4_)_2_HPO_4_, 1.7 g/L citric acid, 0.0025 g/L CoCl_2_, 0.015 g/L MnCl_2_, 0.0015 g/L CuCl_2_, 0.003 g/L H_3_BO_3_, 0.0025 g/L Na_2_MoO_4_, 0.008 g/L Zn(CH_3_COO)_2_, 0.06 g/L Fe(III) citrate, 0.0045 g/L thiamine, 1.3 g/L MgSO_4_, 5 g/L yeast extract and 40 g/L xylose, pH 7.0. In the first stage of the co-culture fermentation, yeast strains were cultured at 28 °C in complex yeast extract peptone dextrose (YPD, all components from BD Biosciences) medium, or synthetic dropout (SD) medium, containing 1.7 g/L yeast nitrogen base (YNB) without amino acids (BD Biosciences), 5 g/L ammonium sulfate (Fisher Scientific), 20 g/L dextrose, and suitable SD mixture at the concentrations specified by the manufacturer (Clontech). XY medium was used in the second stage of the co-culture fermentation. Unless specified, all the chemicals used in this study were purchased from Fisher Scientific or Sigma-Aldrich Co.

### General techniques for DNA manipulation

Plasmid DNA was prepared using the Econospin columns (Epoch Life Science) according to manufacturer’s protocols. PCR reactions were performed using Q5 DNA polymerase (NEB) and Expand High Fidelity PCR System (Roche Life Science) according to manufacturer’s protocols. PCR products were purified by Zymoclean Gel DNA Recovery Kit (Zymo Research). All DNA constructs were confirmed through DNA sequencing by Source Bioscience (LA, USA) or BioBasic (Singapore). Restriction enzymes (NEB) and T4 ligase (NEB) were used to produce and ligate the DNA fragments, respectively. BP Clonase II Enzyme Mix, Gateway pDONR221 Vector and LR Clonase II Enzyme Mix (Invitrogen) and the *S. cerevisiae* Advanced Gateway Destination Vector Kit (Addgene) were used to perform Gateway Cloning (*69*). Using this method, the yeast *expression cassette* vectors were constructed. Plasmids were introduced into yeast cells using Frozen-EZ Yeast Transformation II Kit (Zymo Research). Gibson one-pot, isothermal DNA assembly was conducted at 10 μL scale by incubating T5 exonuclease (NEB), Phusion polymerase (NEB), Taq ligase (NEB) and 50 ng of each DNA fragment at 50 °C for 1 h to assemble multiple DNA fragments into one circular plasmid (*70*). Integrated yeast strains were constructed through homologous recombination and DNA assembly (*71*). Plasmids and microbial strains utilized in this study are listed in tables S1 and S2, respectively. DNA oligonucleotides were synthesized by Integrated DNA Technologies (IDT) and Life Technologies. The plant gene sequences were codon-optimized for expression in *S. cerevisiae* and synthesized by IDT (Coralville, IA) and Twist Bioscience (San Francisco, CA). DNA sequences of genes involved in this work are listed in table S6.

For the construction of *E. coli* expression vectors, the truncated *AtCCD7* gene was amplified by PCR and cloned into the pCDFDuet-1 plasmid (Novagen) using NcoI and NotI to yield the plasmid pCDFDuet-trAtCCD7. The *OsD27* gene was amplified by PCR, digested by NdeI and AvrII and ligated into accordingly digested pCDFDuet-trAtCCD7, yielding the plasmid pCDFDuet-trAtCCD7-OsD27. Using the same strategy, the other six D27 homologous genes were cloned into the pCDFDuet-trAtCCD7 vector, respectively, yielding the corresponding co-expression plasmid. The *OsD27*, truncated *OsD27*, and *AtD27*gene was PCR amplified, digested by NdeI and AvrII and ligated into accordingly digested pCDFDuet-1, yielding the plasmid pCDFDuet-OsD27, pCDFDuet-trOsD27 and pCDFDuet-AtD27. The truncated *AtCCD8* gene was amplified by PCR and cloned into pET21a using Gibson assembly, yielding pET21a-trAtCCD8. The truncated *PpCCD8* was cloned into pET21a, yielding pET21a-trPpCCD8. For the construction of yeast expression cassettes, NADPH-P450 reductase and each individual P450 gene were constructed using Gateway Cloning as described previously. The plasmid of pAC-BETAipi-ispA/dxs was constructed by using Gibson assembly kit (Gibson Assembly® Master Mix, New England Biolabs). The insert was amplified from genomic DNA of *E. coli* BL21(DE3) by using colony PCR and assembled with the backbone of pAC-BETAipi generated by using PCR.

### Culture conditions for *E. coli*-based CL precursors production

For the *in vivo* production of 9CβC, *E. coli* BL21(DE3) was transformed with pAC-BETAipi (Addgene) and pCDFDuet-OsD27, generating *E. coli* **ECL-2**. For 9CβA10′COL production, *E. coli* BL21(DE3) was transformed with pAC-BETAipi (Addgene) and pCDFDuet-OsD27-trAtCCD7 to generate *E. coli* **ECL-3**. Then the yellow colonies were picked and grown in LB with 25 μg/mL chloramphenicol and 50 μg/mL spectinomycin at 37 °C, overnight. 500 µL of the overnight culture was then used to inoculate 5 mL of fresh LB with the corresponding antibiotics with a starting OD_600_ at ∼ 0.05 and cultured at 37 °C and 220 rpm in the 100 mL Erlenmeyer flask. When OD_600_ reached ∼0.6, isopropyl β-D-1-thiogalactopyranoside (IPTG) was added to make the final concentration at 0.2 mM, with ferrous sulfate supplemented at the same time (final concentration at 10 mg/L). Then the cultures were cultivated at 22 °C and 220 rpm for 72 hours.

### Culture conditions for *E. coli*-yeast consortium-based SL production

For the *in vivo* production of SLs, *E. coli* BL21(DE3) was co-transformed with the plasmids pAC-BETAipi (Addgene), pCDFDuet-OsD27-trAtCCD7, pET21a-trAtCCD8, generating *E. coli* **ECL-4**. Single yellow colony was then picked and grown overnight at 37 °C in 1 mL of LB supplemented with 100 μg/mL ampicillin, 25 μg/mL chloramphenicol and 50 μg/mL spectinomycin. 500 µL of the overnight culture was then used to inoculate 5 mL of fresh LB with the corresponding antibiotics with a starting OD_600_ at ∼0.05 and cultured at 37 °C and 220 rpm in the 100 mL Erlenmeyer flask. When OD_600_ reached ∼0.6, IPTG was added with the final concentration at 0.2 mM, with ferrous sulfate supplemented at the same time (final concentration at 10 mg/L). Then the cultures were incubated at 22°C and 220 rpm for 15 hours.

In parallel to preparing the *E. coli* culture, single colony of each yeast strain harboring the corresponding cytochrome P450-expression constructs was used to inoculate an appropriate SD medium considering the auxotrophic markers for maintaining the plasmid(s). The seed culture was incubated at 28 °C and 220 rpm overnight. 100 µL of the overnight grown seed culture was used to inoculate 5 mL of the corresponding SD medium in a 100 mL Erlenmeyer flask and grown at 28 °C for 15 hours.

The *E. coli* and yeast cells prepared as described above were harvested by centrifugation at 3,500 rpm for 5 min. Then the *E. coli* and *S. cerevisiae* cells were mixed and resuspended in 5 mL of XY media (OD_600_ ∼ 8.0) and cultured in 100 mL shake flask at 22 °C and 220 rpm for 60 or 72 hours (final OD_600_ ∼ 40). In the case of CL production, the parent *S. cerevisiae strain (CEN*.*PK2**-**1D)* was pre-cultivated in YPD and mixed with the CL-producing *E. coli* cells.

### Isolation and characterization of SLs and their precursors

Unless specified, 5 mL culture was used for compound extraction. For the extraction of intracellular and extracellular metabolites, 5 mL of cell culture was transferred into 50 mL centrifuge tube, and the cells and medium were separated by centrifugation at 5,000 rpm for 10 min.

The cell pellets were transferred to a new 2 mL microcentrifuge tube and resuspended in 150 μL of dimethylformamide (DMF) and shaken vigorously, followed by the addition of 850 μL of acetone and vigorous shaking for 15 minutes by using a vortex mixer (Thermo scientific), and centrifugation at 12,000 rpm for 10 min. Then the supernatant was transferred to a new 1.7 mL microcentrifuge tube, and dried in a vacuum concentrator (*Eppendorf vacufuge plus*) at 30 °C for 2 to 3 hours. The dried sample was dissolved in 100 μL of acetone.

The medium was transferred into a 50 mL centrifuge tube containing 4 mL of ethyl acetate. The mixture was vortexed vigorously for 5 min by using a vortex mixer (Genie), and then centrifugated at 4,000 rpm for 20 min. The upper ethyl acetate layer of extracted medium was transferred into a 1.7 mL microcentrifuge tube and evaporated to dryness by using a vacuum concentrator (*Eppendorf vacufuge plus*) at 30 °C for 30 minutes. The dried extract was dissolved in 100 μL of acetone.

The samples were centrifuged at 12,000 rpm for 10 min before applied to high performance liquid chromatography analysis. Both UV-VIS and mass spectrometry (MS) detectors were used. The used instrument was Shimadzu LC-MS 2020 (Kyoto, Japan) or a Waters UPLC (ACQUITY) coupled with a Bruker Q-TOF MS (micrOTOF II).

ATβC and 9CβC were analyzed based on *Separation Method I* and a C_18_ column (Kinetex® C18, 100 mm × 2.1mm, 100Å, particle size: 2.6 μm; Phenomex, Torrance, CA, USA). *Separation Method I*: column temperature: 25 °C; single mobile phase: 0.1% (v/v) formic acid in methanol; isocratic elution at 0.4 mL/min; analysis time: 20 min. The injection volume was 10 μL and the UV-VIS absorption was monitored in the range of 190-800 nm. With *Separation Method I*, the retention time of 9CβC was 7.78min, and that of ATβC was 8.17min (characteristic absorption wavelengths: 447 nm and 471 nm for both carotenes).

9CβA10′COL was analyzed using *Separation Method II* and a C_18_ column (Poroshell 120 EC-C18, 100 mm × 3.0 mm, 100Å, particle size 2.7 μm; Aglient, Santa Clara, CA, USA). *Separation Method II:* column temperature: 40 °C; mobile phase A: 0.1% (v/v) formic acid in water; mobile phase B: 0.1% (v/v) formic acid in methanol; gradient elution at 0.5 mL/min. The gradient was as follows: 0–18 min, 5%–100% B; 18–43 min, 100% B; 43–45 min, 100%–5% B. The injection volume was 10 μL and the UV-VIS absorption was monitored in the range of 190-800 nm. With *Separation Method II*, the retention time of 9CβA10′COL was 12.64 min (characteristic absorption wavelengths: 373 nm and 390 nm).

All the SLs (CL, CLA, putative 18-hydroxy-CLA, 4DO, 5DS and OB) were analyzed using *Separation Method III* and a C_18_ column (Kinetex® C18, 100 mm × 2.1 mm, 100Å, particle size 2.6 μm; Phenomex, Torrance, CA, USA). *Separation Method III*: column temperature: 40 °C; mobile phase A: 0.1% (v/v) formic acid in water; mobile phase B: 0.1% (v/v) formic acid in acetonitrile; gradient elution at 0.4 mL/min. The gradient was as follows: 0–28 min, 5%–100% B; 28–35min, 100% B; 35–40min, 5% B. The injection volume was 10 μL and the UV-VIS absorption was monitored in the range of 190-800 nm. With *Separation Method III*, the retention time and the characteristic absorption wavelength of various analytes are listed as follows: βI, 15.91 min (298 nm); CL, 20.58 min (269 nm); CLA, 16.32 min (271 nm), 18-hydroxy-CLA, 12.16 min, 4DO, 15.81 min, OB, 11.03 min, 5DS, 15.97 min. The compounds without the wavelength information were detected using a MS detector, which operates in the m/z range of 50–800 in the positive or negative ion modes. Electrospray Ionization (ESI) was used. The desolvation line temperature was 250 °C. The nebulizing gas and drying gas flow rates were 1.5 L/min and 15 L/min, respectively.

High resolution mass spectrometry (HR-MS) analysis of CLA was performed by using a Ultra Performance Liquid Chromatography (UPLC, Waters ACQUITY) linked with a Time-of-flight Mass Spectrometry (Bruker micrOTOF II). The separation was based on *Separation Method IV* and a C18 column (Poroshell 120 EC-C18 column, 2.1 × 50 mm, particle size 2.7 μm, Agilent Technologies). *Separation Method IV*: column temperature: 40 °C; mobile phase A: 0.1% (v/v) formic acid in water; mobile phase B: 0.1% (v/v) formic acid in acetonitrile; gradient elution at 0.3 mL/min. The gradient was as follows: 0-15 min, 20%-100% B; 15-20 min, 100%-20% B; 20-22 min, 20% B. The MS/MS analysis was performed in product ion scan mode based on ESI (negative mode). The optimized MS/MS conditions were as follows: capillary voltage, 3500 V; end plate offset, 500 V; desolvation gas flow rate (N_2_), 4.0 L/min; drying gas temperature, 200°C; hexapole RF, 50 Vpp; ion energy, 4 eV; collision energy, 18 eV; precursor ion m/z: 331.2. The scan range of m/z was from 50 to 1300. Sodium formate (10 mM) solution was used to calibrate the MS before every use. HR-MS analysis of 18-hydroxy-CLA was performed on a Synapt G2-Si quadrupole time-of-flight mass spectrometer (Waters) coupled to an I-class UPLC system (Waters). The separation was conducted using *Separation Method III* and the C_18_ column (Kinetex® C18, 100 mm × 2.1 mm, 100Å, particle size 2.6 μm; Phenomex, Torrance, CA, USA) as mentioned above. The injection volume was 5 µL. The mass spectra were obtained using the negative ion mode, the scan range of m/z was from 50 to 1200 with a 0.2 s scan time. MS/MS was acquired in a continuum data format and data-dependent fashion with collision energy 25 eV. Source and desolvation temperatures were 150 °C and 600 °C, respectively. Desolvation gas was set to 1100 L/hour and cone gas to 150 L/hour. All gases were nitrogen except the collision gas, which was argon. Capillary voltage was 1.5 kV in negative ion mode.

In the experiment aiming to improve 5DS biosynthesis, β-ionone was analyzed without drying to avoid evaporation loss. 0.5 mL of cell culture was transferred into a 2 mL Eppendorf Safe-Lock Tubes containing 0.1 g glass beads (Sigma, G8772) and 0.5 mL of ethyl acetate. The mixture was incubated at 25 °C and 1,500 rpm for one hour by using a ThermoMixer (Eppendorf), and then centrifuged at 14,000 rpm for 10 min. 100 μL of the upper organic phase was used for Gas Chromatography-Mass Spectrometry (GC-MS, 5977B GC/MSD, Agilent Technologies) analysis. 5 μL of the filtered sample was injected in a spitless mode. HP-5MS capillary column (30 m × 0.25 mm, 0.25 μm film thickness, Agilent Technologies) was used, with helium as the carrier gas at the flow rate of 1 mL/min. The following oven temperature program was carried out: 50 °C for 1 min, 50-100 °C at a rate of 5 °C /min, 100-300 °C at a rate of 50 °C /min, and 300 °C for 1 min. Commercially available βI (Sigma, I12603) was used to prepare standard solutions. The retention time was 11.02 min.

In the experiment aiming to improve 5DS biosynthesis, 5DS was quantified by using the same procedure as the HR-MS experiment, except the used mode was scan instead of product ion scan. No collision energy was applied. The quantification was based on extracted ion chromatogram (m/z: 331.20±0.01). The retention time was 7.95 min.

In the experiment aiming to improve 5DS biosynthesis, the quantification of ATβC and 9CβC was done by using *Separation Method V* and a C30 column (YMC Carotenoid, 250 × 4.6 mm, 5 µm). *Separation Method V*: column temperature: 30 °C; single mobile phase: 75 % (v/v) ethanol, 20 % (v/v) methanol and 5 % (v/v) tetrahydrofuran; isocratic elution at 1 mL/min for 20 min. The injection volume was 10 μL, and the used detector was a UV-VIS detector (475 nm). The model of the HPLC was Agilent 1260 Infinity. The retention time of ATβC and 9CβC were 12.26 min and 13.98 min, respectively.

In the experiment aiming to improve 5DS biosynthesis, the quantification of CL was done using *Separation Method III* on Agilent 1260 Infinity HPLC with a UV detector. The column was Poroshell 120 EC-C18 (150 mm × 4.6 mm, 100Å, particle size 4 μm; Agilent Technologies). The injection volume was 10 μL. The UV detector with a wavelength of 269 nm was used. The retention time of CL was 31.72 min.

## Acknowledgements

pAC-BETAipi was a gift from Francis X Cunningham Jr (Addgene plasmid # 53277; http://n2t.net/addgene:53277;RRID:Addgene_53277). We thank the Metabolomics Core Facility at UC Riverside and Dr. Anil Bhatia for instrument access, training, and data analysis. We thank T.C. for the valuable feedback in the preparation of the manuscript. This work is supported by Cancer Research Coordinating Committee Research Award (grant to Y.L., CRN-20-634571), NIH Director’s New Innovator Award (grant to Y.L., DP2-AT011445), and National Research Foundation Singapore (project IDs: R-279-000-587-592 [through DiSTAP program] and R-279-000-512-281, grants to K.Z.).

## Author contributions

S.W., K.Z. and Y.L. conceived the project; S.W., X.M., A.Z., K.Z. and Y.L. designed the experiments; S.W., X.M., A.Z., and A.V. performed the experiments and analyzed the results; S.W., X.M., A.Z., K.Z., and Y.L. wrote the manuscript.

## Additional information

Competing financial interests: Y.L. and S.W. filed a provisional patent application on Jan. 28, 2021, “Strigolactone-Producing Microbes and Methods of Making and Using the Same,” U.S. Provisional Application No. 63/142,801.

